# Deep learning models detect neuromuscular disease and quantify functional impairment from video-derived biomechanics data

**DOI:** 10.64898/2026.09.30.754736

**Authors:** Sydney C. Covitz, Parker S. Ruth, Shelby Vogt-Domke, Carmichael Ong, Tian Tan, Audrey Chun, Sarah Ismail, Lin Karman, Julie Muccini, Shannon Li, Melina Rogers, Scott Uhlrich, John W. Day, Jennifer L. Hicks, Constance de Monts, Tina Duong, Scott L. Delp

**Author notes:** Corresponding Authors: Sydney C. Covitz & Scott L. Delp.

## Abstract

Measurements of human movement are critical for evaluating patients with neuromuscular diseases. However, current clinical tools are limited in sensitivity and can fail to detect the full spectrum of movement changes patients may exhibit. These limitations can obscure early signs of decline, delaying interventions that could preserve function and independence. OpenCap––a smartphone application that performs video-based markerless motion capture––can help address this gap by enabling scalable, out-of-lab measurements of human movement. Here, we present a method for both detecting disease and quantifying the severity of impaired movement from smartphone videos of a patient’s motion. We used OpenCap to measure kinematics from 337 individuals with diagnoses spanning 12 rare, genetically-defined neuromuscular diseases and 78 age-matched healthy controls. Our dataset is the largest existing publicly available resource of body segment kinematics for this class of diseases. These data were collected in neuromuscular clinics during regular clinic visits as well as at conferences and community events across the United States. We trained deep learning models on these kinematic time series to differentiate between participants with neuromuscular disease and controls (AUROC = 0.97 [0.93-1.00]). We also developed a movement impairment severity score that correlates with both participants’ ratings of their own functional abilities (ρ= -0.81, p<0.001) and existing clinical tests of movement capabilities such as run speed (ρ= -0.83, p<0.001). Our methods for detecting disease and deriving quantitative biomarkers of movement impairment from smartphone videos have the potential to complement clinical assessments, inform screening protocols, and improve clinical trial endpoints.

## INTRODUCTION

Evaluating human movement is a primary means of identifying neuromuscular disease (NMD) and grading functional impairment in patients with movement disorders.^1^ Without routine access to expert clinicians trained to recognize subtle movement abnormalities, patients with rare diseases often experience prolonged diagnostic delays and miss opportunities for early interventions.^2^ Diagnostic delays of several years are especially common in muscular dystrophies, a group of inherited neuromuscular diseases characterized by progressive muscle weakness and diverse onset presentations.^3,4,5^ In the case of myotonic dystrophy (DM), most individuals first seek care from non-neurological specialists and are often misdiagnosed.^5,6^ Diagnostic delays are also common in Charcot–Marie–Tooth disease (CMT), a hereditary peripheral neuropathy with diverse clinical presentations that non-specialists often fail to recognize.^7^ These challenges underscore the need for an approach that can identify movement impairments in settings where expertise and resources are limited.

In-clinic assessment of neuromuscular disease often relies on timed function tests or ordinal, clinician-graded scales that target isolated muscle groups.^8–10^ Timed function tests (e.g., 10-meter gait, timed up-and-go, and sit-to-stand tests) are reproducible, continuous, and quantitative,^11–13^ but these assessments do not capture compensatory changes and patterns of movement quality that may be more indicative of disease than time-based task performance. Strength testing with dynamometry can isolate weakness patterns in specific muscle groups^14^ but provides limited insight into how impairments may affect whole-body functional movements such as gait. Standardized clinical assessments such as the Motor Function Measure (MFM32) provide structured evaluation of whole body motor function and have been validated for tracking disease-specific compensations in gross and fine motor skills.^15^ However, these assessments rely on ordinal scales and lack objectivity, as their administration requires a clinician trained in standardized evaluation methods and compensatory mechanisms typically seen in each disease. Additionally, while researchers have developed high-resolution tools and methods to quantify movement such as optical motion capture and force plates,^16,17^ most clinics lack access to these sophisticated and expensive systems. Further, measuring motion using these tools requires hours of expert labor and trained operators.^18^ Consequently, there remains a need for scalable, objective measures of whole-body function across heterogeneous impairment presentations that can be obtained from data acquired in neuromuscular clinics.

Recent advances in video-based movement analysis offer an objective and accessible way to quantify full body motion. Smartphone-based tools such as OpenCap^19^ enable markerless motion capture using commodity hardware, making quantitative biomechanical data collection feasible both inside and outside the clinic. Prior work by our team showed that kinematic features derived from OpenCap videos can capture disease-specific movement signatures in DM and facioscapulohumeral muscular dystrophy (FSHD).^20^ While these expert-specified features were able to distinguish between two types of muscular dystrophy, the ability for video-derived biomechanics to capture the full range of movement patterns that arise in a larger sample of neuromuscular diseases with heterogeneous presentations remains untested.

Deep learning models trained on full movement trajectories, rather than a small set of discrete features, have the potential to interpret and quantify a wide range of impairments. These models can analyze long sequences by learning patterns associated with disease directly from the kinematics without engineering prior knowledge into the training process.^21,22^ Such models applied to video data of full body motion have demonstrated strong performance in predicting clinical gait parameters in children with cerebral palsy.^23^ However, their application in rare neuromuscular diseases has been limited by the availability of large-scale, high-quality movement data.

To address this limitation, we collected, curated, and publicly released the Neuromuscular OpenCap Dataset,^24^ which comprises 675 visits from 415 participants with 12 different neuromuscular disorders and healthy controls (see **Table 1**), with the largest representations from DM, FSHD, and CMT. Using OpenCap, we recorded our participants performing a 15-minute protocol that included up to nine standardized tasks to evaluate gait, balance, and upper-extremity function. Participants range in age from 4-84 years (mean ± SD: 42.6 ± 18.1 years, see **Table 2, Sup. Fig. 1**). These data were collected across the United States in clinical, conference, community event settings. Our cohort includes recordings our team had previously collected and released^25^ and has been widely expanded to add 10 new rare diseases and 281 new participants with a range of movement capabilities. The size and diversity of this unique movement dataset enable us to leverage modern deep learning model architectures^21^ with the capacity to learn patterns from complex, heterogeneous inputs.

**Fig. 1.**
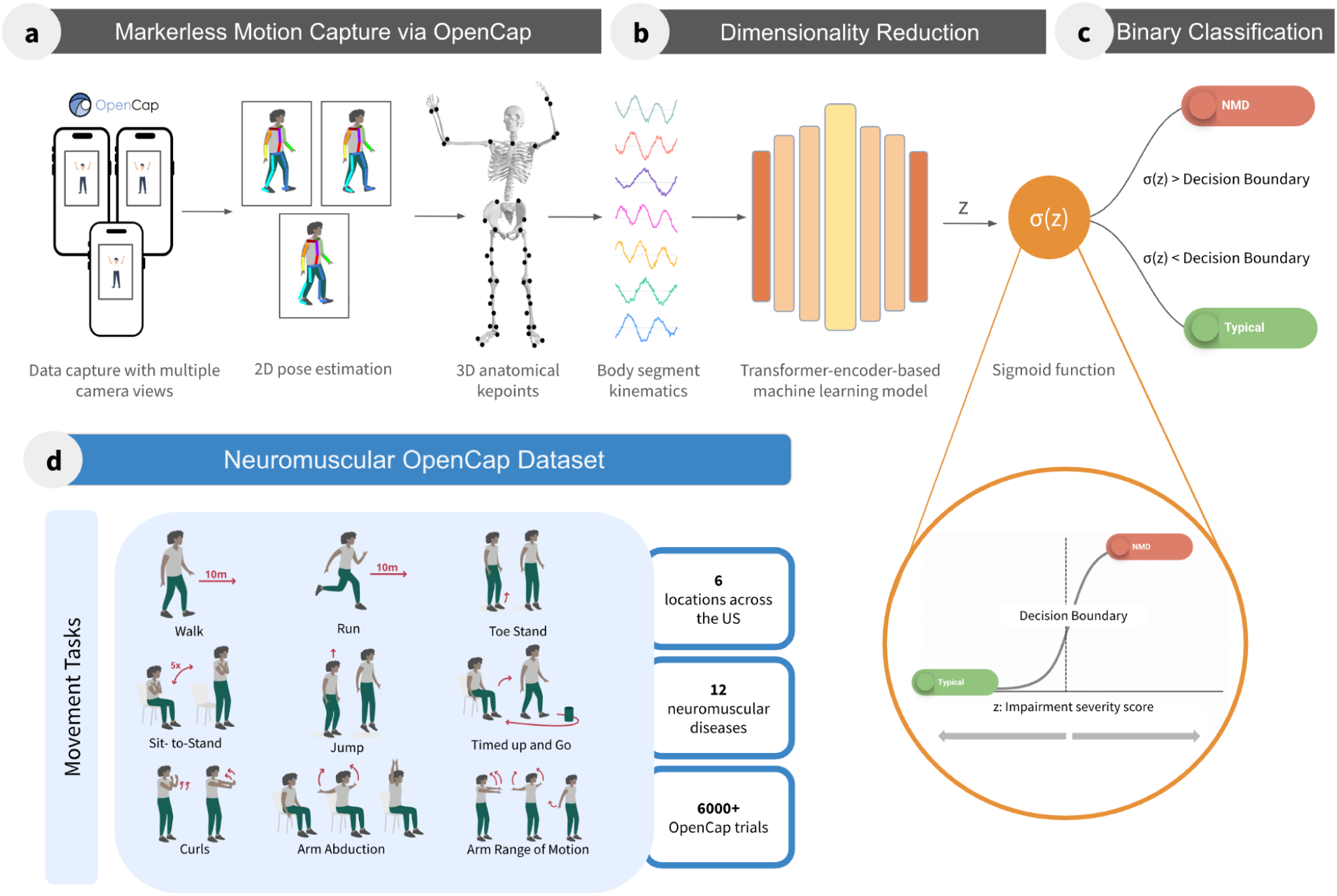
| Video-based disease detection and impairment severity scoring. (a) Videos of movement tasks were captured with OpenCap. A binary classification model, comprising (b) a machine learning model that features a transformer encoder block and (c) a sigmoid function for classification, was trained to detect the presence of a neuromuscular disease. Model inputs are time series of body segment kinematics derived from OpenCap videos of movement. The trained model’s pre-sigmoid output (z) is a continuous movement impairment severity score. Our model was trained on the Neuromuscular OpenCap Dataset (d).

**Table 1.** | Participant and visit counts by diagnostic group in the Neuromuscular OpenCap Dataset. Each visit corresponds to one data collection session. Female percentage is reported at the participant level. NMD = neuromuscular disease, CTL = healthy control.

| Group | Participants<br>(n) | Female<br>n (%) | Visits<br>(n) |
| --- | --- | --- | --- |
| **NMD** | **337** | **166 (49.3%)** | **535** |
| Myotonic dystrophy (DM) | 138 | 86 (62.3%) | 248 |
| Facioscapulohumeral muscular dystrophy (FSHD) | 54 | 14 (25.9%) | 110 |
| Charcot-Marie-Tooth disease (CMT) | 77 | 46 (59.7%) | 86 |
| Duchenne muscular dystrophy (DMD) | 20 | 0 (0.0%) | 26 |
| Spinal muscular atrophy (SMA) | 9 | 4 (44.4%) | 14 |
| Becker muscular dystrophy (BMD) | 8 | 0 (0.0%) | 10 |
| Cystinosis myopathy (CM) | 9 | 7 (77.8%) | 16 |
| Amyotrophic lateral sclerosis (ALS) | 2 | 2 (100.0%) | 2 |
| GNE myopathy (GNE) | 10 | 2 (20.0%) | 12 |
| Limb-girdle muscular dystrophy (LGMD) | 3 | 1 (33.3%) | 4 |
| Spinobulbar muscular atrophy (SBMA) | 3 | 0 (0.0%) | 3 |
| Symptomatic carrier of DMD (scDMD) | 4 | 4 (100.0%) | 4 |
| **CTL** | **78** | **41 (52.6%)** | **140** |
| **Total (NMD + CTL)** | **415** | **207 (49.9%)** | **675** |

**Table 2.** | Visit-level demographics and functional measure spreads. Demographic and clinical functional measures across the Neuromuscular OpenCap Dataset cohort, split into neuromuscular disease (NMD) and healthy control (CTL) groups. Each visit corresponds to one data collection session. Demographics (age, height, weight) are reported as mean ± SD. Functional measures (10m run and walk times, TUG, 5×STS, Brooke upper-extremity score, ACTIVLIM sum) are reported as median (range). The Brooke score is a clinician-graded ordinal measure of upper extremity function that uses a 1-6 scale; a score of 6 indicates typical arm abduction ability, and lower values indicate greater impairment.

| Metric | NMD | CTL | Total |
| --- | --- | --- | --- |
| Demographic measures |  |  |  |
| Age (years) | 43.9 ± 17.7 | 37.5 ± 19.1 | 42.6 ± 18.1 |
| Height (m) | 1.7 ± 0.1 | 1.6 ± 0.2 | 1.7 ± 0.2 |
| Weight (kg) | 71.8 ± 19.9 | 64.8 ± 21.8 | 70.4 ± 20.5 |
| Functional measures |  |  |  |
| 10 m run time (s) | 6.0 (2.8–24.4) | 3.2 (2.4–4.9) | 4.8 (2.4–24.4) |
| 10 m walk time (s) | 9.5 (5.8–42.5) | 7.6 (4.8–13.4) | 9.0 (4.8–42.5) |
| TUG time (s) | 8.0 (4.3–42.6) | 5.7 (4.2–8.8) | 7.2 (4.2–42.6) |
| 5×STS time (s) | 11.9 (5.2–45.6) | 7.2 (3.9–17.8) | 10.8 (3.9–45.6) |
| Brooke score | 6.0 (2.0–6.0) | 6.0 (5.0–6.0) | 6.0 (2.0–6.0) |
| ACTIVLIM Score | 2.7 (-7.1–7.2) | 7.2 (4.3–7.2) | 3.5 (-7.1–7.2) |

This paper introduces a data-driven method for detecting and quantifying movement impairment in patients with neuromuscular diseases from videos of movement tasks that can be performed in neuromuscular clinics (**Fig. 1**). Our goals were twofold: (1) detect the presence of neuromuscular disease–related movement impairment from video-derived kinematics, and (2) estimate the severity of an individual’s functional impairment on a continuous scale. We trained machine learning models on our Neuromuscular OpenCap Dataset to predict the presence of neuromuscular disease from kinematic time series extracted from smartphone videos of participants performing movement tasks. We then derived a continuous movement impairment severity score that quantifies the degree to which an individual’s movement deviates from healthy control reference patterns. Finally, we assessed the clinical relevance of this score by examining its association with timed function tests and patient-reported outcomes.

## RESULTS

### Detecting neuromuscular disease from video

We developed an encoder-only transformer-based model that ingested OpenCap-derived body segment kinematics extracted from videos of movement tasks and learned a decision boundary between NMD patients and healthy controls performing movement tasks. The transformer achieved strong disease detection performance across five-fold cross-validation and a held-out test set. On the pooled out-of-fold (OOF) validation set, each participant visit––one data collection session––was scored by the model from the fold in which it was held out. All visits from a single participant were part of the same fold. Resulting predictions were then pooled across all five folds, achieving an area under the receiver operating characteristic curve (AUROC) of 0.92 [0.90-0.95], area under the precision–recall curve (AUPRC) of 0.98 [0.97-0.99], and balanced accuracy (bACC) of 0.87 [0.83-0.90] (**Fig. 2a**). Bracketed values are 95% confidence intervals obtained by bootstrap resampling over visits. On the held-out test set, each visit was scored by an ensemble of the five cross-validation models (the mean of the five models’ predicted disease probabilities). The held-out test ensemble achieved an AUROC of 0.97 [0.93-1.00], AUPRC of 0.99 [0.98-1.00], and bACC of 0.86 [0.75-0.96] (**Fig. 2a**).

**Fig. 2.**
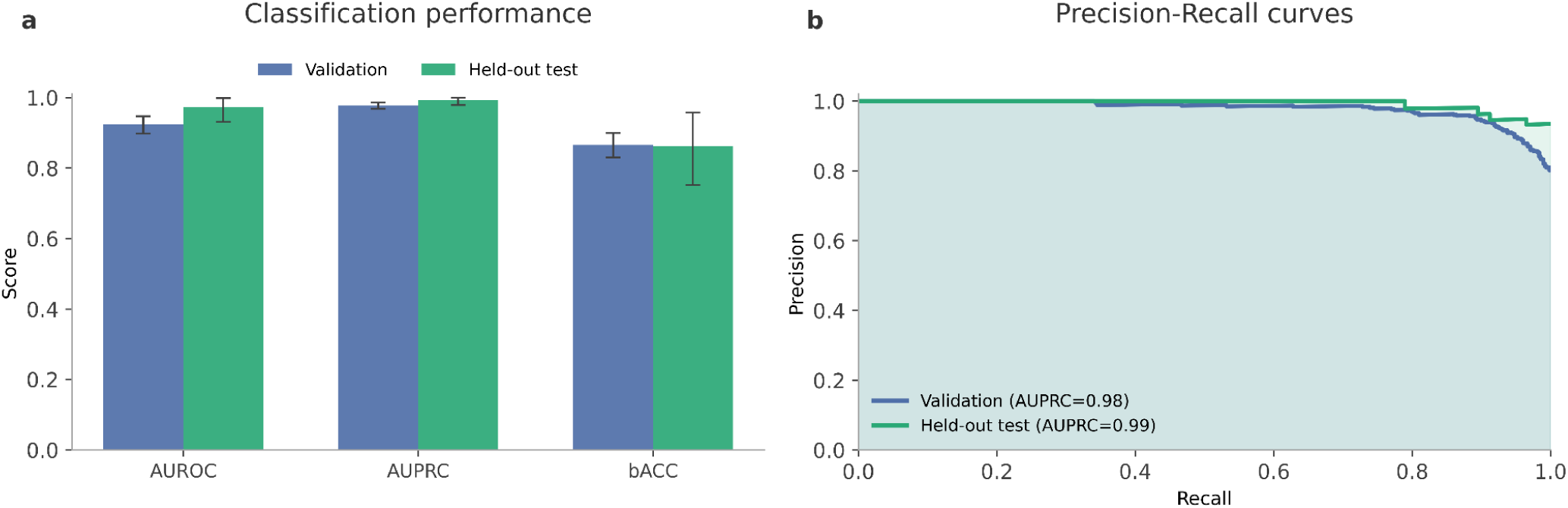
| Performance of the transformer model for disease detection on validation and held-out test sets. (a) Area under the receiver operating characteristic curve (AUROC), area under the precision-recall curve (AUPRC), and balanced accuracy (bACC) for pooled out-of-fold (OOF) validation (blue) and ensemble-averaged held-out test (green) predictions. The held-out test ensemble prediction for each visit is the mean of the disease probabilities predicted by the five trained models from cross-validation. Error bars are 95% confidence intervals from bootstrap resampling over visits. (b) Precision–recall curves for pooled out-of-fold validation predictions (blue) and ensemble-averaged held-out test predictions (green). Shaded regions show the area under each curve (AUPRC) for both OOF validation and held-out test sets.

Precision–recall analysis showed high precision across a broad range of recall for both OOF validation (AUPRC = 0.98) and held-out test (AUPRC = 0.99), with test predictions averaged across the five-fold models closely tracking the OOF curve (**Fig. 2b**). Performance was stable across the five independently trained cross-validation models. Evaluated individually on the held-out test set, they achieved AUROC 0.94 ± 0.02, AUPRC 0.98 ± 0.01, and balanced accuracy 0.88 ± 0.05 (mean ± SD), indicating the result reflects a consistent signal across training splits rather than a single favorable fold. To assess whether disease detection performance was robust to different neural network types, rather than dependent on architectural choice, we additionally trained a support vector machine (SVM) and a multilayer perceptron (MLP) on hand-crafted summary statistics extracted from the same kinematic time series. Detection performance was comparable across model architectures (**Sup. Table 1**), indicating the discriminative signal present in the kinematics is robust to model choice rather than an artifact of a single architecture.

### Computing the movement impairment severity score

Having established that OpenCap-derived kinematics contain a strong disease detection signal, we investigated whether that signal also carries information about the degree of movement impairment across individuals. We defined our movement impairment severity score as the continuous output of the transformer encoder (z in **Figure 1**), prior to applying the sigmoid function and thresholding for binary classification (**Figures 1, 7**). For the held-out test data, the severity score was computed from the mean value of z across all five cross-validation fold models, yielding a stable ensemble estimate. NMD participants had significantly higher severity scores than healthy controls (p < 0.001). Higher scores indicate greater deviation from typical movement. Statistical significance was assessed with a two-sided Mann–Whitney U test, and Cliff’s δ is the accompanying effect size (95% bootstrap confidence interval). The severity score showed strong separation between healthy controls and participants with neuromuscular diseases in the held-out test set, with Cliff’s δ = 0.92, 95% CI [0.82, 0.99], indicating that a randomly selected NMD participant had a higher severity score than a randomly selected healthy control in 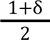 = 96% of pairings (**Fig. 3a-b**). Out-of-fold cross-validation results showed comparable separation between NMD participants and controls (Cliff’s δ = 0.85, 95% CI [0.80, 0.89], **Fig. 3c-d**). Scores varied substantially across NMD diagnoses (**Fig. 3b, d**), reflecting the range of functional abilities present within our cohort (see **Table 2, Sup. Fig. 1**).

**Fig. 3.**
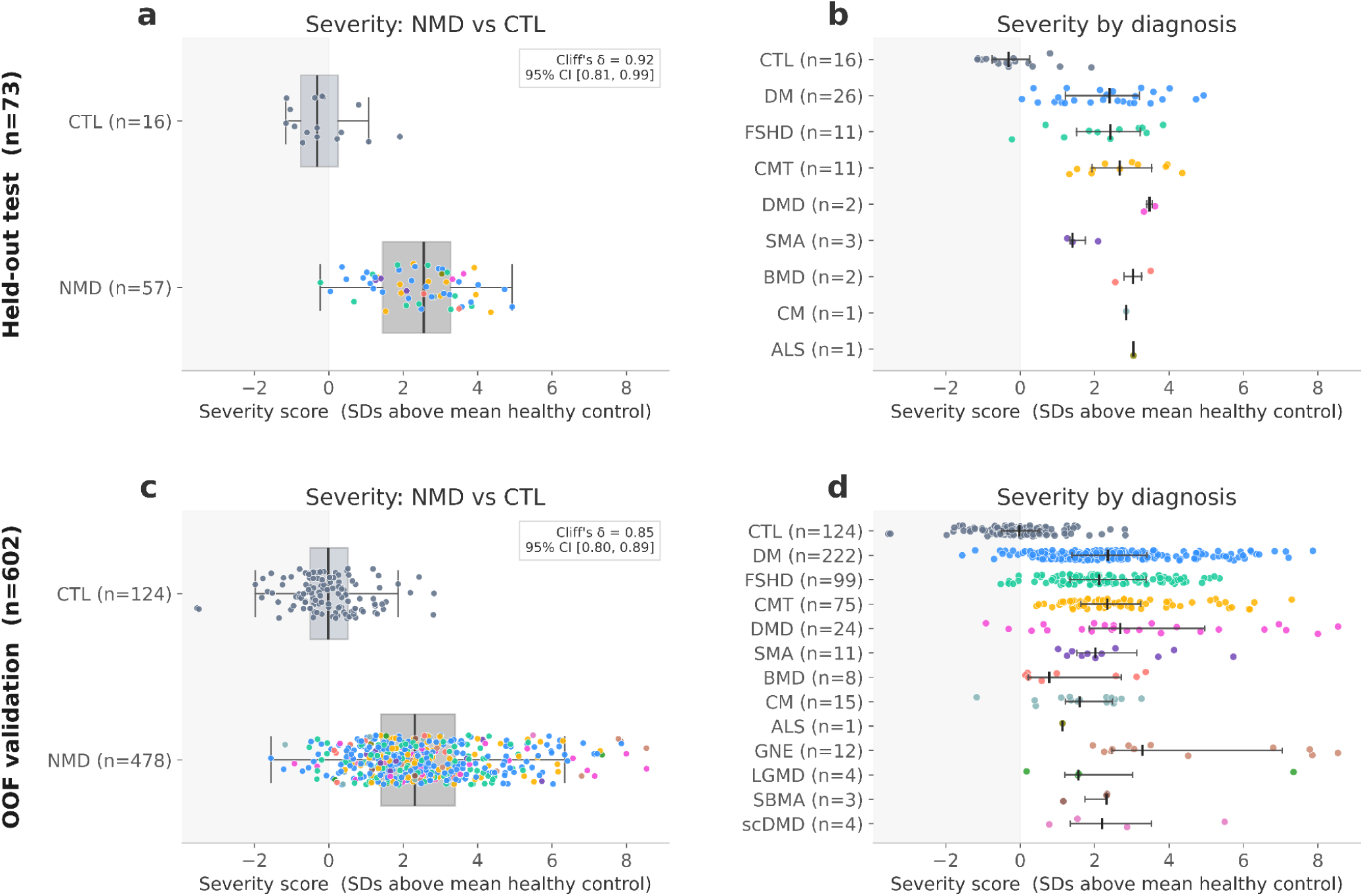
| Movement impairment severity score distribution on validation and held-out test sets. Movement impairment severity scores for held-out test set visits (a) and OOF validation (c) are grouped as healthy control (CTL) vs. neuromuscular disease (NMD). Each point represents one visit, and colors indicate clinical diagnosis. Black vertical lines indicate the median severity score for each group, and boxes show the interquartile range (IQR) with whiskers extending to the most extreme values within 1.5 times the IQR of the quartiles. Sample sizes are shown in parentheses. Scores were normalized so that zero reflects the mean severity score of the out-of-fold (OOF) validation controls, with one unit corresponding to one standard deviation of control scores. Effect size between disease and control groups was quantified using Cliff’s δ. The same severity scores are stratified by clinical diagnosis for held-out test data (b) and OOF validation (d). Each point represents one visit, the vertical line marks the median, and the horizontal line indicates the interquartile range. DM = myotonic dystrophy, FSHD = facioscapulohumeral muscular dystrophy, CMT = Charcot-Marie-Tooth disease, DMD = Duchenne muscular dystrophy, SMA = spinal muscular atrophy, BMD = Becker muscular dystrophy, CM = cystinosis myopathy, ALS = amyotrophic lateral sclerosis, GNE = GNE myopathy, LGMD = limb-girdle muscular dystrophy, SBMA = spinobulbar muscular atrophy, scDMD = symptomatic carrier of Duchenne muscular dystrophy.

### Validating the movement impairment severity score

Our movement impairment severity score correlated with patient-reported survey responses from the ACTIVLIM^26^ questionnaire, which evaluates ability to perform everyday functional tasks and has been validated for patients with neuromuscular disorders (**Fig. 4**). The severity score achieved strong correlation with ACTIVLIM across the held-out test set (ρ = −0.81, p < 0.001), and a moderate correlation when restricting to neuromuscular disease only (ρ = −0.62, p < 0.001). Our severity score also correlated with timed function tests commonly conducted in clinic (**Fig. 5**), with higher scores corresponding to slower task performance (10m run speed: ρ = −0.83, p < 0.001; timed up and go time: ρ = 0.73, p < 0.001; 10m walk speed: ρ = −0.63, p < 0.001, and 5×STS time: ρ = 0.56, p < 0.001).

**Fig. 4.**
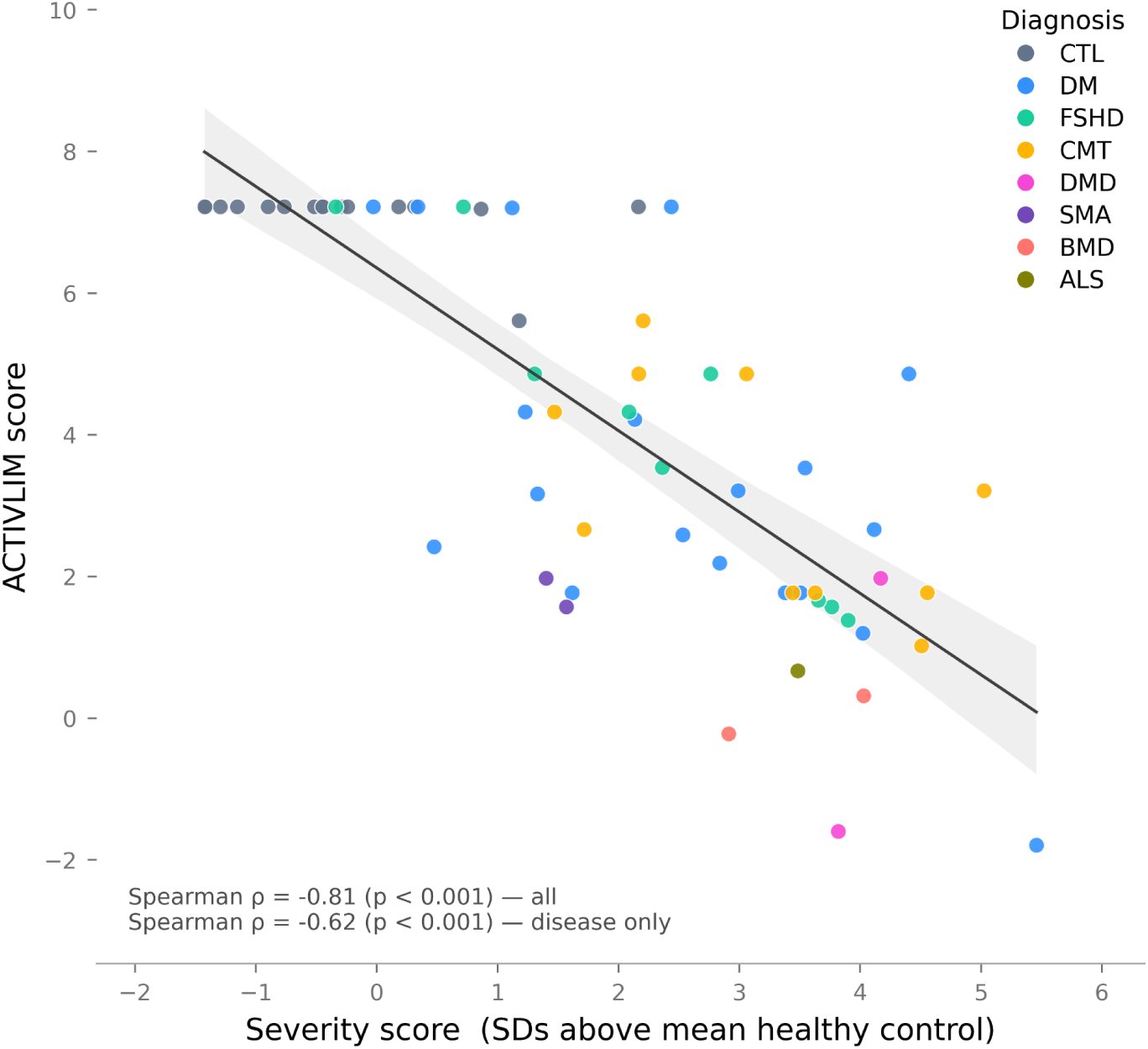
| Validation of the movement impairment severity score against patient-reported measures of function. Patient-reported function was quantified as a Rasch logit computed from the 22 questions on the ACTIVLIM questionnaire, which measures self-reported difficulty of everyday functional tasks on a three-point ordinal scale (0 = impossible, 2 = easy). Only held-out test set visits where the participant completed the survey are shown (n=61 total, n=45 with a neuromuscular disease). The solid line is a linear regression fit across all visits, and the shaded band is its 95% bootstrap confidence interval. Spearman’s ρ and p-values are shown both for all visits and for neuromuscular disease only.

**Fig. 5.**
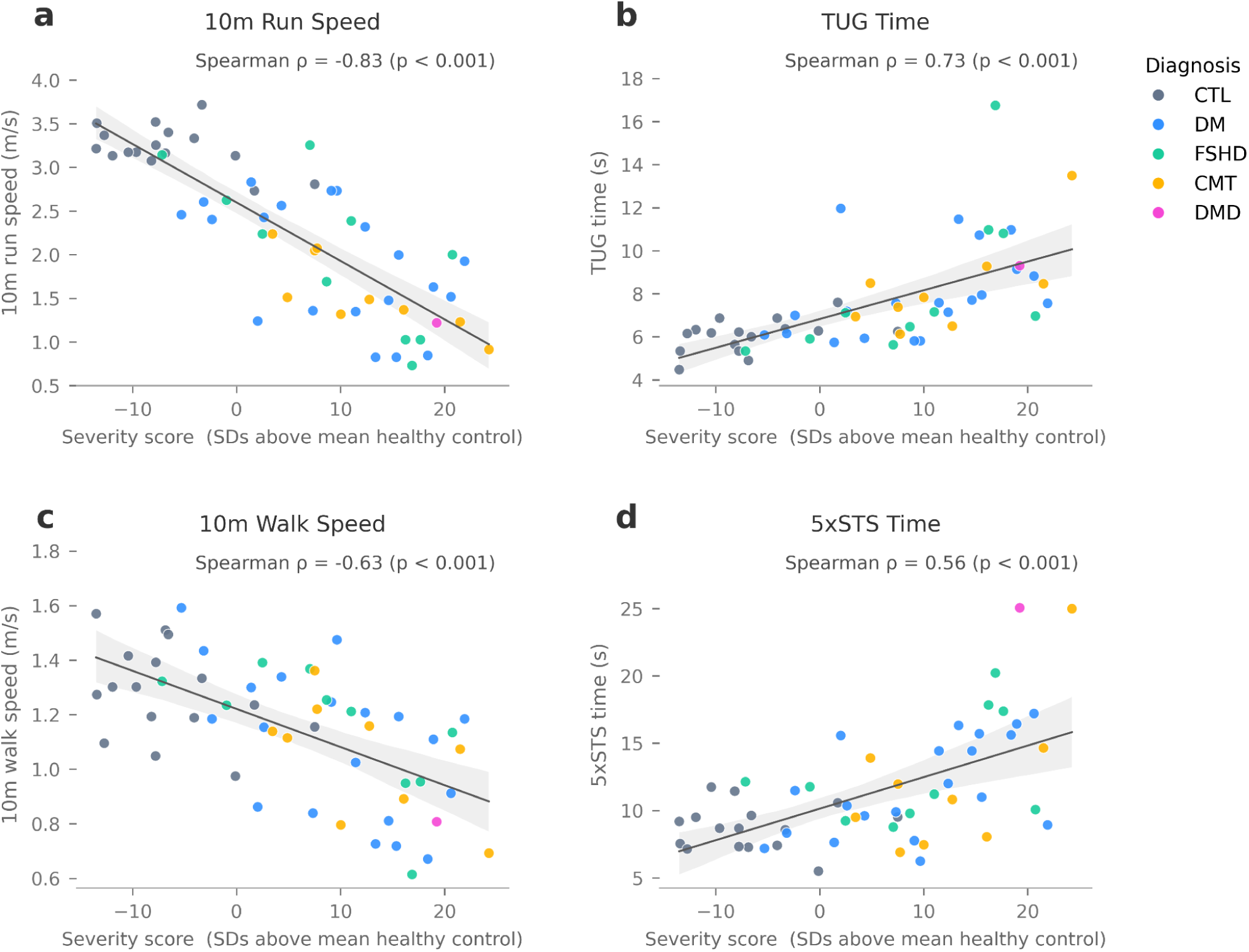
| Validation of the movement impairment severity score against clinical measures of function. Correlations between the movement impairment severity score and (a) 10m run speed, (b) timed up-and-go (TUG) time, (c) 10m walk speed, and (d) five times sit-to-stand (5×STS) time in the held-out test set. Only held-out test set visits without missing data for any of the four timed function tests are shown (n=56 visits). The solid line is a linear regression fit across all visits, and the shaded band is its 95% bootstrap confidence interval. Spearman’s ρ and p-values are shown per panel.

## DISCUSSION

We developed and validated a method for detecting and quantifying neuromuscular disease using kinematic time series derived from smartphone video recordings of a 15-minute movement protocol. We achieved high performance on disease detection with an AUROC of 0.97 across the eight distinct diseases in our held-out test set (**Fig. 2**). Our video-based classifier is not intended to replace existing practices for detecting diseases, which involve in-clinic evaluations followed by confirmatory genetic testing and muscle biopsies.^27^ Instead, our disease detection results suggest that kinematics-based methods could augment existing protocols by flagging disease-indicating movement patterns and accelerating patients gaining access to the specific genetic panel they require. Beyond binary detection, we derived a continuous movement impairment severity score (**Fig. 3**) from the model’s pre-sigmoid output that achieved high separation between controls and diseased participants (Cliff’s δ = 0.92, **Fig. 3a**). Our score’s correlation with participants’ ratings of their own functional abilities (ρ= -0.81, **Fig. 4**) and existing clinical movement tests such as run speed (ρ= -0.83, **Fig. 5a**) provides evidence that our method captures functional consequences. Unlike measures that are separately engineered^20^ or reliant on ordinal scales,^10,15,28,29^ our severity score emerges from the detection objective itself. Since it is anchored to the distribution of healthy movement rather than to a disease-specific instrument, our score places every participant regardless of diagnosis on a common impairment axis. This could help inform physical therapy protocols and enable the selection of comparable cohorts for natural history studies and clinical trials.

Prior work in movement-based disease detection has largely focused on identifying a single disorder or estimating severity within one diagnosis.^30^ For example, smartphone accelerometry has distinguished children with Duchenne muscular dystrophy from typically developing peers using gait features, with classification accuracy reaching 100% in selected task-specific analyses.^31^ In Parkinson’s disease detection, machine learning classifiers using a single lumbar sensor achieved a test AUROC of 0.85,^32^ while a model leveraging multi-sensor inputs achieved an AUROC of 0.98.^33^ Models trained on multi-sensor wearable data have also discriminated between Parkinson’s disease motor stages with AUROCs of 0.76-0.90.^34^ In contrast, our model distinguished participants with neuromuscular disease from controls with an AUROC of 0.97 in a held-out test set that included eight different neuromuscular disorders using body segment kinematics derived from smartphone video. This places our held-out test-set performance within the range reported for high-performing movement-based classifiers, while extending the level of discrimination to a clinically heterogeneous group of neuromuscular disorders using data collected from commodity hardware (iPhones).

Digital movement biomarkers of disease severity obtained from single task recordings have been associated with patient-reported functional outcomes in single-condition cohorts. In FSHD, a kinematics-based composite score derived from an instrumented Timed Up and Go correlated with PROMIS Physical function survey results (r = -0.58).^35^ Kneel-to-stand time extracted from Microsoft Kinect recordings of patients after lower-extremity sarcoma treatment correlated with the Toronto Extremity Salvage Score (r = -0.56).^36^ Our severity score’s correlations with ACTIVLIM (ρ = -0.81, ρ = -0.62 for NMD only) are similar in magnitude to existing associations between single-task digital movement measures and patient experiences of physical function, suggesting that a multi-task, cross-disease score can retain clinically meaningful relationships with functional limitation across neuromuscular disorders.

An important limitation of our work is that participants were not stratified by disease onset or duration. Our dataset captures a range of severity—evidenced by the spread of scores within each disease subpopulation (**Fig. 3**)—but we cannot make claims about performance in early or prodromal disease. A critical next step is evaluation in patients with confirmed early-stage disease when the benefit of detection would be greatest. Additionally, our cohort is subject to selection bias. Because we recruited many participants at disease-specific advocacy conferences (including the Myotonic Dystrophy Foundation, FSHD Society, Hereditary Neuropathy Foundation (HNF), and the Charcot-Marie-Tooth Association (CMTA), our sample likely skews toward people already engaged with specialty care and patient communities. Performance may therefore differ in primary care settings, where the target population is less selected, and validating the model in those settings is an important next step.

Several directions would extend the current pipeline toward greater clinical utility. Our current model collapses all neuromuscular diseases into a single disease label; future disease-specific classifiers trained within this framework could add differential diagnosis capabilities alongside detection. Additionally, there are multiple distinct genetic subtypes of DM (2), FSHD (2) and CMT (>80). We hope the continued growth of the Neuromuscular OpenCap Dataset will aid our ability to define features and patterns common to each genetic form. Further, decomposing the whole-body score into segment-specific sub-scores for the upper limb, lower limb, trunk, and pelvis could provide clinically actionable localization of impairment to guide rehabilitation planning and monitor segment-specific disease progression independently of overall severity. Finally, all our analyses were purely cross-sectional, and we have not yet established the score’s sensitivity to longitudinal change nor its test-retest reliability. Whether our severity score is responsive to change over time—the prerequisite for use as a trial endpoint—is therefore an important question for future work. We anticipate that the public release of our dataset, models, and analysis code will advance the field in these directions.

Despite the central role of motor function in neuromuscular disease, objective, quantitative methods to detect and grade it in the clinic remain limited. Most established methods are disease-specific, ordinal, and depend on clinician administration and judgment,^9,15,28,29^ which constrains comparison across diseases and examiners. This gap is especially consequential because several of the neuromuscular diseases represented in our dataset have no approved disease-modifying therapies yet are the focus of active clinical trials; the lack of sensitive, objective outcome measures remains an ongoing barrier to trial success.^27,37^ Our continuous movement impairment severity score, which quantifies whole-body function is a first step towards addressing this need. Building on prior digital biomarker studies,^20,38–41^ our approach uses an end-to-end model trained on body segment kinematics extracted from smartphone videos of movement to detect and quantify impairment across 12 neuromuscular diseases without disease-specific feature engineering. Because our detection and severity quantification method requires only smartphones and a brief movement protocol that can be administered in a clinic, it may provide a scalable approach for incorporating quantitative neuromuscular disease screening and tracking into clinical research and practice.

## METHODS

### Data collection using OpenCap

We collected video recordings of movement tasks using OpenCap which estimates body segment kinematics from smartphone videos. Individuals performed up to nine movement tasks they could safely complete. These tasks span gait, strength, balance, and upper- and lower-extremity function. We cleaned, curated, and released these data publicly on Zenodo as the Neuromuscular OpenCap Dataset.^24^ This dataset contains 675 visits from 415 unique participants, including individuals with 12 rare, genetically-defined neuromuscular diseases and healthy controls (**Table 1**, **Table 2, Sup. Fig. 1**). Our cohort is primarily ambulatory with the exception of 23 self-reported nonambulatory participants. We define non-ambulatory as a participant who was unable to perform 10m gait tasks. Data were collected across eight US states in clinical, conference, and community event settings. We used DataLad—a version control software package for scientific data management—to ensure reproducibility and provenance tracking throughout the data curation process.^42^

The data collection setup and protocol followed those in Ruth et al.^20^ During each visit, participants were instructed to perform up to nine activities: gait at a maximal, self-selected speed, timed up-and-go, sit-to-stand, jump, toe-stand, shoulder abduction, arm range-of-motion, and bicep curls. Participants skipped tasks that they or their clinicians deemed unsafe to perform. All participants provided written informed consent to the collection, processing, and optional sharing of their data under a protocol approved by the Stanford University Institutional Review Board (IRB; protocols IRB-23888 and IRB-39283). The OpenCap system is HIPAA-compliant and has been authorized by the Stanford University Offices of Privacy and Data Security to process and store identifiable personal health information.

### Model architectures and training

We trained an encoder-only transformer to detect and quantify neuromuscular disease from OpenCap-derived kinematics. We define disease detection in this context as the ability to identify individuals with a confirmed diagnosis compared to those without one from human movement kinematics. We also trained two other model types (a support vector machine and a multilayer perceptron) on the same cross-validation/held-out test sets splits to assess the robustness and stability of the kinematic signal across architectures of varying complexity.

#### Input preparation

Kinematic time series were extracted from smartphone video recordings using OpenCap, which fits a musculoskeletal model using OpenSim to estimate body segment kinematics from synchronized, multi-view smartphone video. These kinematics span degrees of freedom of the pelvis, lower limbs, lumbar spine, and upper limbs and were used as inputs to our machine learning models (**Sup. Table 3**). All trials were conducted without assistive devices, and all sessions were processed with the HRNet^43^ pose detector, v0.3 of OpenCap’s marker enhancer^44^, and the LaiUhlrich2022^19^ musculoskeletal model. Our data release^24^ includes estimated marker positions, kinematics, scaled models, and metadata dictionaries (one for each data collection session) detailing the camera information and OpenCap settings used to collect and process that session’s trials.

We held out 10% of the data as a test set. The remaining 90% was divided into five-fold cross-validation subsets with no participant appearing in more than one split (GroupKFold^45^). Within each fold, 80% of the data were used for training and 20% for validation. Time series were smoothed using a zero-lag 4th order Butterworth low-pass filter with a cutoff frequency of 6 Hz and normalized per kinematic parameter using the mean and standard deviation computed from each fold’s training split only. These normalized statistics were applied without recomputation to the validation and test sets.

#### Encoder-only transformer

Our primary model is an encoder-only transformer trained end-to-end on the kinematic time series extracted from OpenCap (**Fig. 6**). Visits contain up to nine movement tasks, each represented as a set of time series (one for each of the 33 body segment kinematic parameters).

**Fig. 6.**
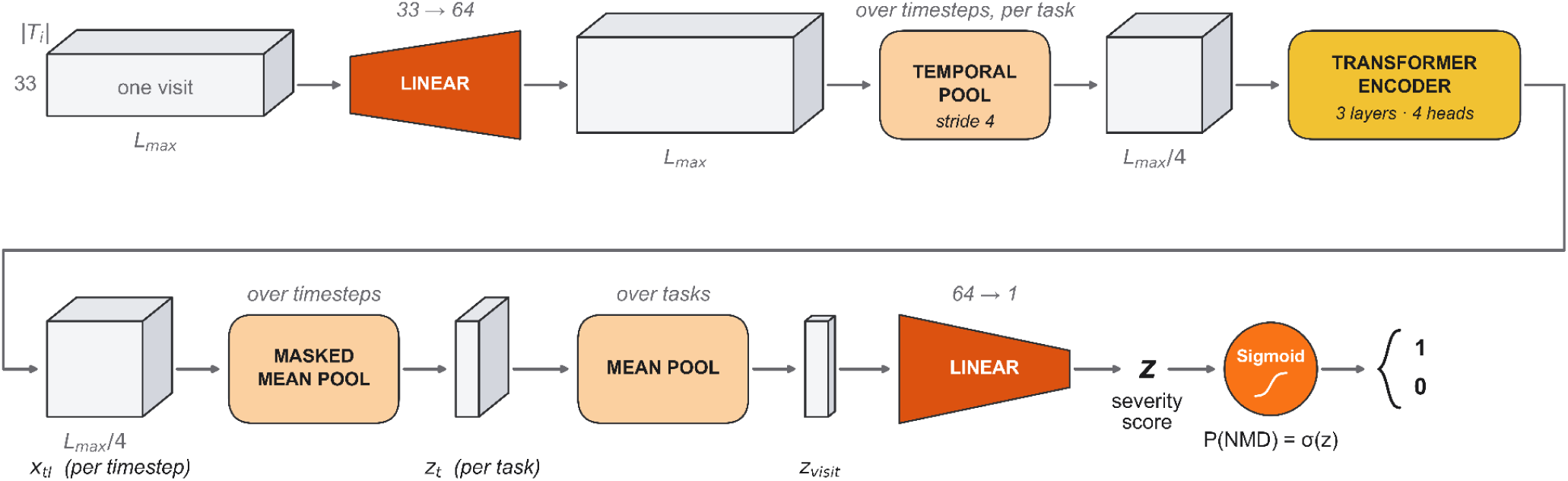
| Path of a single visit passed through the transformer-encoder-based model architecture for neuromuscular disease detection and severity scoring. Grey blocks represent data tensor shapes. Orange 2D shapes represent neural network layers. Each movement task in a visit is encoded independently, then collapsed over its valid, non-padded timesteps into a single task embedding. Task embeddings are averaged across all tasks in a visit to form one visit-level representation, which a linear layer maps to a scalar (the movement impairment severity score). A sigmoid converts this score to a probability of neuromuscular disease, which a decision threshold then classifies as either disease or control.

Within each mini-batch of four visits, all task sequences across visits were stacked into a single tensor of shape [T_batch_, L_max_, 33], where T_batch_ is the total number of tasks across visits in the batch, L_max_ is the maximum sequence length in the batch, and 33 is the number of body segment kinematic parameters whose trajectories we use as model inputs (**Sup. Table 3**). Sequences were zero-padded to the length of the longest sequence in the batch, with a validity mask indicating which timesteps contained kinematic data versus padding.

For each task, a linear projection layer first mapped the 33-dimensional kinematic time series to a 64-dimensional space in order to grant the transformer attention heads more room to encode different aspects of each movement pattern while also satisfying the technical requirement that our model dimensions must be divisible by the number of attention heads (4). To reduce sequence length prior to attention, temporal average pooling was applied over non-padded timesteps with stride 4 (non-overlapping windows), averaging only the valid frames in each window. A pooled window is retained as valid when more than half of its frames are non-padding. The pooled sequence is processed by a three-layer transformer encoder with the same validity mask passed to each attention layer to prevent attention over padded positions.

After encoding, a task-level representation *z_t_* is obtained by averaging encoder outputs over non-padded timesteps only:

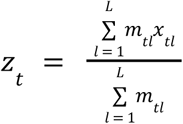

where *x_tl_* ∈ ℝ^64^ is the encoder output at timestep *l* for task *t* and *m_tl_* ∈ {0, 1} is the validity mask that indicates whether timestep *_l_* contains kinematic data. Task-level embeddings are then averaged across all tasks completed in the same visit:

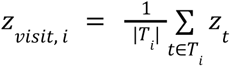

Where *T_i_* is the set of tasks belonging to visit *i*, |*T_i_* | is the number of tasks completed in that visit, and *z_t_* ∈ ℝ^64^ is the task-level embedding. This yields one 64-dimensional visit-level representation regardless of how many tasks are present, accommodating missing tasks without requiring special handling. This visit-level representation is then passed through a sigmoid classification head to produce a disease probability in (0,1). A threshold of 0.5 is then applied to that probability to obtain the disease vs. healthy prediction.

#### Support vector machine and multilayer perceptron

To assess whether disease detection performance is robust across degrees of model complexity rather than dependent on architectural choice, we additionally trained a support vector machine (SVM) and a multilayer perceptron (MLP) on hand-crafted summary statistics extracted from our same kinematic time series. Specifically, we computed five features per kinematic parameter—mean, standard deviation, range, RMS velocity, interquartile range––as well as per task mean absolute acceleration. Mean absolute acceleration was computed as the mean absolute second finite difference of each kinematic time series, averaged across 33 body segment parameters, providing a scalar measure of movement smoothness per task. All task-level feature vectors were aggregated to the visit level via max pooling across tasks, selecting the maximum value of each feature across all tasks performed in a given visit.

#### Model training and validation

All models were trained using five-fold grouped cross-validation (GroupKFold by participant). The transformer was optimized with AdamW^46^ (learning rate=1e-3, weight decay=1e-4). After each epoch, the model was evaluated on that fold’s validation split, and the checkpoint with the highest validation AUROC was retained. Training stopped early if validation AUROC did not improve for five consecutive epochs.

### The best model checkpoint per fold was saved based on maximum validation AUROC. We employed class-weighted binary cross-entropy loss

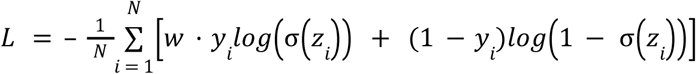

where 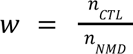 = is the positive class weight computed from the training split of each fold, *y_i_* ∈ {0, 1} is the binary disease label for visit *i* (neuromuscular disease = 1, healthy control = 0), *_zi_* is the pre-sigmoid logit, and *_N_* is the number of visits in the batch. The global L2 norm of all model gradients was clipped at each optimization step.

### Disease detection performance

Disease detection performance was evaluated using area under the receiver operating characteristic (AUROC), area under the precision-recall curve (AUPRC), and balanced accuracy (bACC) across all three models’ cross-validated classification results. Because the cohort is class-imbalanced, we report metrics that are robust to prevalence rather than raw accuracy. We summarize discrimination with AUROC, which is threshold-free and prevalence-independent, and AUPRC, which reflects performance on the minority (control) class. Balanced accuracy––the mean of sensitivity and specificity––was computed at a fixed probability threshold of 0.5, applied identically to the out-of-fold (OOF) validation and held-out test sets. We report performance as mean ± 95% bootstrap confidence intervals across five folds for both OOF validation and held-out test sets. Confidence intervals were obtained by bootstrap resampling over visits.

### Movement impairment severity score construction

The transformer’s pre-sigmoid logit—the scalar value produced by the final linear layer prior to the sigmoid activation function (**Fig. 7**)—was used as a continuous movement impairment severity score. For the SVM and MLP, we computed an analogous severity score using the log-odds transform of the predicted disease probability,

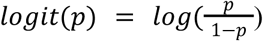

which maps calibrated probabilities from (0,1) back onto the real number line, placing all three models’ scores on a common unbounded scale. For the held-out test set, severity scores were computed as the mean logit across all five trained fold models, yielding an ensemble estimate per participant. For interpretability purposes, severity scores were normalized so that zero corresponds to the mean logit of healthy controls in the cross-validation set, and one unit corresponds to one standard deviation of that distribution. On this scale, a score of zero reflects the average healthy controls, and higher scores reflect greater deviation from typical movement and increasingly characteristic of neuromuscular disease.

### Movement impairment severity score validation

To assess whether our movement impairment severity score captures clinically meaningful variation in impairment status, we correlated our score with four timed function test metrics—10m run speed, 10m walk speed, 5xSTS time, and TUG-cone time—and the ACTIVLIM patient-reported outcome measure, a survey that assesses difficulty with activities of daily living.^26^ Spearman’s rank correlations were computed across all test set participants with available outcome data.

## DATA AVAILABILITY

Neuromuscular OpenCap Dataset––the motion dataset we collected and curated for this work––is freely available for download on Zenodo at the following link: https://doi.org/10.5281/zenodo.22309771.

## CODE AVAILABILITY

We have publicly released all trained models along with the code we used to train and inference them. Our GitHub repository can be found at the following link: https://github.com/stanfordnmbl/NMD_Detection_Quantification. We have also released an example walkthrough (see our [README.md](https://dealsbe.com/) file) that provides instructions users can follow to reproduce each result reported in this manuscript.

## ACKNOWLEDGEMENTS

We thank all our volunteers including Shirley Paulose, Tia Nguyen, Veronica Schnyer, Lesly Welsh, Whitney Tang, Sally Dunaway Young, Libby Yatsu, Rabia Farooquee, Lisa Ghiglieri, Raiye Hailu, Nathan Hageman, Meng Yao, Paxton Ataide, Sabrina Salvatore, Sanchalee Khonde, Meghan Moore, Carolyn Kelley, Melissa McIntyre, and others from Stanford University and Children’s Hospital of Colorado who helped facilitate data collection. We especially wish to thank our 415 study participants who made this research possible. We are also grateful to the Myotonic Dystrophy Foundation, the FSHD Society, and the Charcot-Marie-Tooth Association (CMTA) and Hereditary Neuropathy Foundation (HNF) for their support of our study. This work was funded by the Wu Tsai Human Performance Alliance and the Mobilize Center under NIH Grant P41EB027060. The funders played no role in project design, data collection, analysis and interpretation of data, or the writing of this manuscript.

## AUTHOR CONTRIBUTIONS

S.C.C., P.S.R., J.H., T.T., T.D., J.W.D., and S.L.D. conceptualized and designed the project.

C.O., S.U., J.M., C.M., T.D, and S.L.D. developed the methods used to collect data.

S.V.D., S.I., L.K., S.C.C., P.S.R., S.U., T.T., M.R., C.M., and T.D. collected data

S.C.C., S.L., S.V.D., and P.S.R cleaned and processed data

S.C.C. developed the models and generated figures

S.C.C., P.S.R., and S.L.D. drafted and revised the manuscript.

J.H., T.T., C.O., C.M., A.C., S.L, S.V.D., J.M., J.W.D., and T.D. revised the manuscript.

## COMPETING INTERESTS

S.U. is a cofounder of Model Health, Inc., which supports the non-academic, commercial use of OpenCap––the open source software used for data collection in this study.

## Notes

https://zenodo.org/records/22309771

https://github.com/stanfordnmbl/NMD_Detection_Quantification

