## Supplementary Material for "Deep learning models detect neuromuscular disease and quantify functional impairment from video-derived biomechanics data"

Distribution of visit-level demographic and functional measures

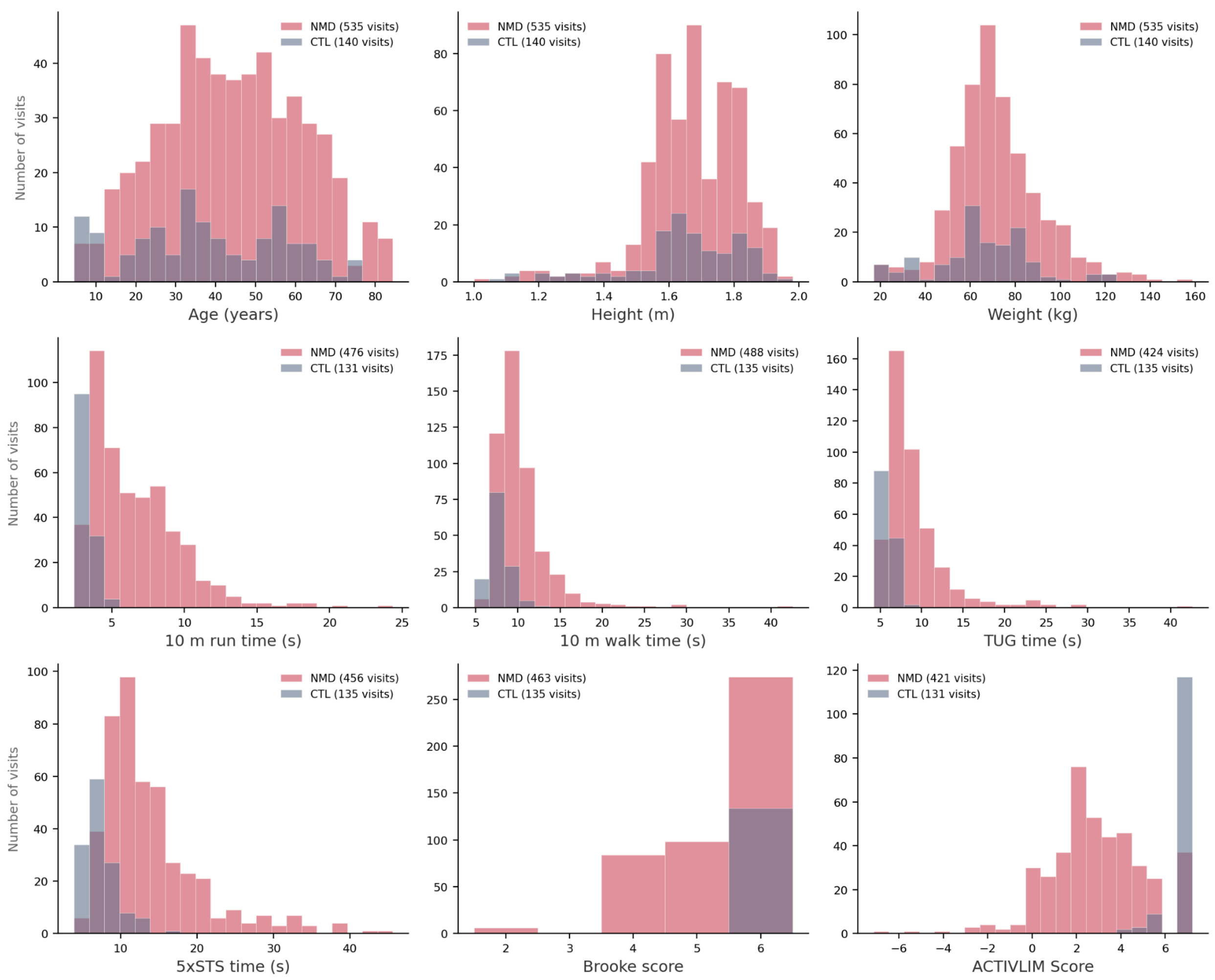


**Supplementary Fig. 1 | Distribution of demographic and functional measures.** Visit-level histograms of the nine measures summarized in Table 2, for neuromuscular-disease (NMD) and control (CTL) groups. Every visit is a data point, and the y-axis is the number of visits. The number of visits in the bottom 6 subplots varies, as not all participants did all tasks at each visit. The panels show the spread and group overlap behind Table 2's summary statistics.

| Section | Measure | Transformer | SVM | MLP |
|---|---|---|---|---|
| Ensemble test-set performance [95% CI] | AUROC | 0.97 [0.93, 1.00] | 0.96 [0.90, 0.99] | 0.95 [0.88, 0.99] |
| | AUPRC | 0.99 [0.98, 1.00] | 0.99 [0.97, 1.00] | 0.98 [0.96, 1.00] |
| | bACC | 0.86 [0.75, 0.96] | 0.80 [0.68, 0.92] | 0.85 [0.73, 0.95] |
| Difference vs Transformer [95% CI] | AUROC | ref. | −0.02 [−0.07, +0.02] | −0.03 [−0.08, +0.02] |
| | AUPRC | ref. | −0.00 [−0.02, +0.01] | −0.01 [−0.03, +0.00] |
| | bACC | ref. | −0.06 [−0.19, +0.08] | −0.01 [−0.13, +0.11] |

**Supplementary Table 1 | Disease detection performance by model architecture.** Three different model architectures were trained on the same recordings and evaluated on the same held-out test set (73 visits from 42 participants; 57 NMD, 16 typical). The transformer operates on kinematic time series. The SVM and MLP operate on summary statistics aggregated for each visit. Differences were computed on the five model ensemble prediction, e.g. Δ = AUROC(SVM ensemble) − AUROC(Transformer ensemble), and assessed by a paired bootstrap over visits (5,000 resamples). Both models were evaluated on the same resampled visits so that their correlated errors cancel. No difference from the transformer was statistically significant (all 95% CIs included 0). bACC was evaluated at threshold=0.5.

| Section | Measure | Transformer | SVM | MLP |
|---|---|---|---|---|
| Spearman $\rho$ between clinical measures and severity score | ACTIVLIM ($n = 61$) | 0.81 | 0.68 | 0.69 |
| | TUG-cone ($n = 60$) | 0.71 | 0.70 | 0.64 |
| | 5×STS ($n = 63$) | 0.57 | 0.62 | 0.58 |
| | 10 m walk time ($n = 67$) | 0.58 | 0.57 | 0.53 |
| | 10 m run time ($n = 67$) | 0.77 | 0.77 | 0.72 |
| $\Delta\rho$ vs Transformer [95% CI] | ACTIVLIM | ref. | −0.13 [−0.25, −0.02]* | −0.12 [−0.27, +0.00] |
| | TUG-cone | ref. | −0.02 [−0.14, +0.12] | −0.07 [−0.22, +0.08] |
| | 5×STS | ref. | +0.05 [−0.08, +0.18] | +0.01 [−0.15, +0.15] |
| | 10 m walk time | ref. | −0.01 [−0.15, +0.13] | −0.05 [−0.20, +0.10] |
| | 10 m run time | ref. | −0.00 [−0.10, +0.10] | −0.05 [−0.18, +0.06] |

**Supplementary Table 2 | Spearman correlations between three models' severity scores and clinical measures on the held-out test set.** Clinical measures were sign-aligned so that a higher value indicates worse function, which matches the direction of our movement impairment severity score. Differences are expressed relative to the transformer (negative would indicate a weaker correlation than the transformer) and were estimated by a paired bootstrap over visits, with both models evaluated on each resample. Correlations with the four timed function tests were indistinguishable across model families (no difference reached significance). The transformer correlated more strongly with ACTIVLIM than the SVM (95% CI excluded zero).

| Kinematic parameter | OpenSim coordinate | Side |
|---|---|---|
| **Pelvis** | | |
| Pelvis tilt | `pelvis_tilt` | – |
| Pelvis list | `pelvis_list` | – |
| Pelvis rotation | `pelvis_rotation` | – |
| Pelvis anterior/posterior translation | `pelvis_tx` | – |
| Pelvis vertical translation | `pelvis_ty` | – |
| Pelvis mediolateral translation | `pelvis_tz` | – |
| **Trunk** | | |
| Lumbar extension | `lumbar_extension` | – |
| Lumbar bending | `lumbar_bending` | – |
| Lumbar rotation | `lumbar_rotation` | – |
| **Lower limb** | | |
| Hip flexion | `hip_flexion_{r,l}` | L/R |
| Hip adduction | `hip_adduction_{r,l}` | L/R |
| Hip rotation | `hip_rotation_{r,l}` | L/R |
| Knee flexion | `knee_angle_{r,l}` | L/R |
| Ankle dorsiflexion | `ankle_angle_{r,l}` | L/R |
| Subtalar angle | `subtalar_angle_{r,l}` | L/R |
| MTP angle | `mtp_angle_{r,l}` | L/R |
| **Upper limb** | | |
| Arm flexion | `arm_flex_{r,l}` | L/R |
| Arm adduction | `arm_add_{r,l}` | L/R |
| Arm rotation | `arm_rot_{r,l}` | L/R |
| Elbow flexion | `elbow_flex_{r,l}` | L/R |
| Forearm pronation/supination | `pro_sup_{r,l}` | L/R |

**Supplementary Table 3 | OpenSim body segment kinematics used as disease detection model inputs.** 33 OpenSim body segment kinematic parameters that form the transformer-based model's input feature vector at each timestep, listed by anatomical segment (pelvis, lower limb, trunk, upper limb). Pelvis pose comprises three orientation coordinates (tilt, list, rotation) and three translations (anterior/posterior, vertical, mediolateral). All remaining coordinates are joint angles. Angular coordinates are expressed in degrees, and pelvis translations are expressed in meters. R and L denote the right and left sides. MTP = metatarsophalangeal joint.
